# Fast centromeric repeat turnover provides a glimpse into satellite DNA evolution in *Nothobranchius* annual killifishes

**DOI:** 10.1101/2023.03.23.534043

**Authors:** Anna Voleníková, Karolína Lukšíková, Pablo Mora, Tomáš Pavlica, Marie Altmanová, Jana Štundlová, Šárka Pelikánová, Sergey A. Simanovsky, Marek Jankásek, Martin Reichard, Petr Nguyen, Alexandr Sember

## Abstract

Satellite DNA (satDNA) is rapidly evolving class of tandem repeats with some motifs being involved in centromere organization and function. Rapid co-evolution of centromeric satDNA and associated proteins has been mostly attributed to the so-called centromere drive. To identify repeats associated with centromeric regions and test for the role of meiotic drive in their evolution, we investigated satDNA across Southern and Coastal clades of African annual killifishes of the genus *Nothobranchius*. C-banding showed expansion of (peri)centromeric heterochromatin regions in the Southern-clade killifishes. Molecular cytogenetic and bioinformatic analyses further revealed that two previously identified satellites, Nfu-SatA and Nfu-SatB, are associated with centromeres only in one lineage of the Southern clade. Nfu-SatB was, however, detected outside centromeres also in other members of the Coastal clade, which is consistent with the “library” hypothesis of satDNA evolution. We also identified a novel satDNA, Cl-36, associated with (peri)centromeres in *N. foerschi*, *N. guentheri* and *N. rubripinnis* from the Coastal clade. Our findings could be explained by centromere drive shaping karyotype change and centromeric repeat turnover in *Nothobranchius* species with possible reversal of spindle polarity within the Southern clade.

## 1. Introduction

African killifishes from the genus *Nothobranchius* (Peters, 1868) (Aplocheiloidei: Nothobranchiidae) are small freshwater fishes with bigger and more colorful males compared to smaller and dull females (Wildekamp 2004; Berois et al. 2016). The genus is monophyletic and currently comprises over 90 species (Nagy and Watters 2021; Fricke et al. 2023) partitioned into seven evolutionary lineages (van der Merwe et al. 2021). *Nothobranchius* spp. are adapted to periodic droughts driven by cycles of rainy and dry seasons in south-eastern African savannahs, where they survive in isolated pools, temporarily flooded by rainwater (Blažek et al. 2013; Furness 2016; Cellerino et al. 2016). Having the shortest life cycle among vertebrates, the turquoise killifish *N. furzeri* (Jubb, 1971) became a popular model system for aging research (Cellerino et al. 2016; Hu and Brunet 2018). Besides, the unique biology of killifishes offers many advantages for studies related to developmental biology, population dynamics and evolution (Cellerino et al. 2016; Terzibasi Tozzini and Cellerino 2020). For instance, their mating system and sexual dimorphism make them attractive for studies of reproductive isolation and sexual selection (Berois et al. 2016; Cellerino et al. 2016).

*Nothobranchius* killifishes became of interest also to genome and sex chromosome research. Studies reported high repetitive DNA content in *Nothobranchius* genomes (Reichwald et al. 2009, 2015; Cui et al. 2019; Štundlová et al. 2022) and wide variation in diploid chromosome numbers (2n = 16–50) and karyotype structures in 73 studied representatives (Krysanov et al. 2016; Krysanov and Demidova 2018; Krysanov et al. 2023). Moreover, a multiple sex chromosome system of the X_1_X_2_Y type has been cytogenetically identified in six distant *Nothobranchius* spp., which suggests remarkable sex chromosome evolution (Ewulonu et al. 1985; Krysanov et al. 2016; Krysanov and Demidova 2018). Intriguingly, the *N. furzeri* genome sequence revealed an XY sex chromosome pair with polymorphic size of a non-recombining region in different populations (Reichwald et al. 2015; Willemsen et al. 2020). It was hypothesized that the *N. furzeri* Y chromosome polymorphism represents an early stage of sex chromosome evolution (Reichwald et al. 2015). However, physical mapping of various repeats in *N. furzeri* and its sister species *N. kadleci* revealed that repetitive DNA landscape differs considerably between their X and Y chromosomes and these differences extend beyond the non-recombining regions. One particular difference between the X and Y chromosomes was a largely reduced block of constitutive heterochromatin in the centromeric region on Y chromosomes in two out of three examined populations (Štundlová et al. 2022). This region overlapped with hybridization signals of fluorescence *in situ* hybridization (FISH) with two repeats associated with *N. furzeri* centromeres, the Nfu-SatA and Nfu-SatB (Reichwald et al. 2009; Štundlová et al. 2022).

Certain satellite DNAs (satDNA), i.e. tandemly repeated DNA class with rapid molecular evolution (Plohl et al. 2012; Garrido-Ramos 2017; Thakur et al. 2021), can be associated with centromeres (Melters et al. 2013; Hartley and O’Neill 2019; Talbert and Henikoff 2020) and thus are considered to be involved in segregation of chromosomes during cell divisions (Henikoff et al. 2001; McKinley and Cheesman 2016). Yet despite their rather conservative function, centromeric satDNAs turn over very fast (Henikoff et al. 2001; Bracewell et al. 2019; Ávila Robledillo et al. 2020; Nishihara et al. 2021). It has been hypothesized that rapid co-evolution of both centromeric DNA and associated proteins is mainly driven by centromere drive (Henikoff et al. 2001). The hypothesis postulates that homologous chromosomes differ in their capability to bind spindle microtubules and thus can segregate non-randomly exploiting the asymmetric female meiosis, which produces three polar bodies (i.e. the evolutionary dead-ends) and only one egg (Henikoff et al. 2001; Kursel and Malik 2018; Kumon and Lampson 2022). Deleterious segregation errors induce selective pressure, which fuels the evolution of involved proteins and DNA repeats, thus suppressing the drive (Henikoff et al. 2001; Kumon and Lampson 2022).

Hence, it was hypothesized that the reduction in centromeric clusters of the Nfu-SatA and Nfu-SatB repeats on Y chromosomes observed in *N. furzeri* reflects relaxed selection imposed by centromere drive (Štundlová et al. 2022), as the Y chromosome never passes through female meiosis (cf. Yoshida and Kitano 2012; Pokorná et al. 2014). Unfortunately, nothing is known about killifish centromeric organization outside *N. furzeri* and *N. kadleci* (Reichwald et al. 2009, 2015; Štundlová et al. 2022) and little is known about the centromere organization in teleost fishes in general. Rather than identifying sequences which bind centromeric proteins (Cech and Peichel 2016; Ichikawa et al. 2017), the available studies have focused mainly on sequences associated with centromeres, detected either by molecular or bioinformatic methods and physically mapped by means of *in situ* hybridization (Ferreira et al. 2010; Suntronpong et al. 2020; Stornioli et al. 2021; Goes et al. 2022, 2023; Kretschmer et al. 2022). More recently, these sequences have been inferred directly from long read sequencing data (Ichikawa et al. 2017; Conte et al. 2019; Varadharajan et al 2019; Tao et al. 2021). These are typically satellite sequences presumably containing conserved motifs such as the CENP-B box needed for chromosome stability and cell division (Suntronpong et al. 2016; Gamba and Fachinetti 2020). Centromeric tandem repeats seem to be homogenized at higher rates in teleost fishes compared to other vertebrates (Suntronpong et al. 2020) and their evolutionary dynamics seems to reflect centromeric position with those of acrocentrics being more conserved (Ichikawa et al. 2017).

In the present study, we analyzed repetitive sequences across the representatives of *Nothobrachnius* genus by means of RepeatExplorer2 bioinformatic pipeline (Novák et al. 2020) to identify repeats associated with centromeric regions and to look for evidence of centromere drive. Our results suggest that Nfu-SatA and Nfu-SatB are associated with centromeres only in one lineage of the Southern clade, although Nfu-SatB can be detected also in representatives of the Coastal clade, in agreement with the “library” hypothesis (i.e. the existence of shared collection of satDNA repeats among related species, with varied degree of their amplification and contraction; Fry and Salser 1977; Ruiz-Ruano et al. 2016). We also identified novel repeat associated with centromeres in the Coastal-clade species. Based on the presence of larger (peri)centromeric heterochromatin blocks observed in the Southern-clade species but not in other studied representatives, we discuss a possible reversal of spindle orientation in the common ancestor of this clade.

## 2. Materials and Methods

### 2.1 Fish sampling

We analyzed individuals of 14 species representing the Southern and Coastal clade (seven and five species, respectively) of the genus *Nothobranchius*, with *N. ocellatus* and *Fundulosoma thierryi* as their outgroups. The studied individuals from *N. orthonotus*, *N. kuhntae*, *N. pienaari*, *N. rachovii*, *N. eggersi* and *N. rubripinnis* were sampled from laboratory populations recently derived from wild-caught individuals and were previously identified based on morphology and the phylogenetic analysis of mitochondrial and nuclear DNA markers (for details, see Bartáková et al. 2015; Blažek et al. 2017; Reichard et al. 2022). The remaining species were obtained from specialists and experienced hobby breeders who keep strictly population-specific lineages derived from original imports. In this case, the species identity was confirmed on the basis of key morphological characters (Wildekamp 1996, 2004; Watters et al. 2008, 2020; Nagy 2018). The detailed information is provided in Table 1.

**Table 1.** List of *Nothobranchius* killifish species used in this study along with their sample sizes (N) and origin

| Clade | Species | Code | N* | Source / locality |
| --- | --- | --- | --- | --- |
| outgroup | *Fundulosoma thierryi* Ahl, 1924 | FTH | 1♂, 3♀ | aquarium strain |
| Southern clade | *Nothobranchius furzeri* Jubb, 1971 | NFU | 1♂, 1♀ | Chefu, MZ |
|  | *N. kadleci* Reichard, 2010 | NKA | 1♂, 1♀ | Gorongosa, MZ |
|  | *N. orthonotus* (Peters, 1844) | NOR | 3♂, 3♀ | Limpopo, MZ |
|  | *N. kuhntae* (Ahl, 1926) | NKU | 4♂, 3♀ | Pungwe, MZ |
|  | *N. pienaari* Shidlovskyi, Watters & Wildekamp, 2010 | NPI | 2♂, 3♀ | Limpopo, MZ |
|  | *N. krysanovi* Shidlovskyi, Watters & Wildekamp, 2010 | NKR | 2♂, 2♀ | Quelimane, MZ |
|  | *N. rachovii* Ahl, 1926 | NRA | 2♂, 2♀ | Beira Airport, MZ |
| Ocellatus clade | *N. ocellatus* (Seegers, 1985) | NOC | 1♂, 1♀ | Nyamwage, TZ |
| Coastal clade | *N. eggersi* Seegers, 1982 | NEG | 2♂, 1♀ | Bagamoyo, TZ |
|  | *N. foerschi* Wildekamp & Berkenkamp, 1979 | NFO | 2♂ | Soga, TZ |
|  | *N. guentheri* (Pfeffer, 1983) | NGU | 4♂, 3♀ | Zanzibar, TZ |
|  | *N. cardinalis* Watters, Cooper & Wildekamp, 2008 | NCA | 1♂ | Matandu, TZ |
|  | *N. rubripinnis* Seegers, 1986 | NRU | 2♂, 2♀ | Kitonga, TZ |

\* number and sex of individuals used for each method is specified in Supplementary Table 1

### 2.2 Chromosomal preparations

Mitotic chromosome spreads were obtained either i) from regenerating caudal fin tissue (Völker and Ráb 2015) with modification described in Sember et al. (2015) and a fin regeneration time ranging from one to two weeks, or ii) by a direct preparation from the cephalic kidney following Ráb and Roth (1988) and Kligerman and Bloom (1977), with in the latter protocol being modified according to Krysanov and Demidova (2018). In the kidney-derived preparations, the chromosomal spreading quality was enhanced using a dropping technique by Bertollo et al. (2015). Preparations were inspected with phase-contrast optics and those of sufficient quality were dehydrated in an ethanol series (70%, 80%, and 96%, 2 min each) and stored at -20 °C until use.

### 2.3 Constitutive heterochromatin staining

Analysis of constitutive heterochromatin distribution was done by C-banding (Haaf and Schmid 1984), using 4′,6-diamidino-2-phenolindole (DAPI) (1.5 μg/mL in anti-fade; Cambio, Cambridge, UK) counterstaining. Fluorescent staining with the GC-specific fluorochrome Chromomycin A_3_ (CMA_3_) and the AT-specific fluorochrome DAPI (both Sigma-Aldrich, St. Louis, MO, USA) was performed according to Mayr et al. (1985) and Sola et al. (1992).

### 2.4 Whole-genome sequencing data

Genomic DNA was sequenced *de novo* in *Nothobranchius guentheri*, *N. kadleci*, *N. orthonotus*, *N. rachovii* and *N. rubripinnis*. First, high molecular weight genomic DNA (HMW gDNA) was extracted from three females of each species using a MagAttract HMW DNA Kit (Qiagen, Hilden, Germany), following the provided protocol. Next, Illumina paired-end libraries with 450 bp insert size were prepared from the isolated HMW gDNA and sequenced on the NovaSeq 6000 platform at Novogene (HK) Co., Ltd. (Hong Kong, China), yielding, at least, 5 Gb (ca 3.3× coverage of *Nothobranchius furzeri* genome; 1C = 1.54 Gb, Reichwald et al. 2009). Resulting data were deposited into the Sequence Read Archive (SRA) under accession numbers XXX – XXX. In *N. furzeri*, sequencing data from three female specimen were obtained from the SRA (accession numbers ERR583470, ERR58471, SRR1261480; Reichwald et al. 2015).

### 2.5 Analysis of repetitive DNA

The satellitome was characterized using RepeatExplorer2 (Novák et al. 2020). Prior to the analysis, the quality of raw Illumina reads was checked using FastQC (version 0.11.5; Andrews 2010). Low quality reads and adapter sequences were removed using cutadapt (version 1.15; Martin 2011) with settings for two-color chemistry: ‘--nextseq-trim=20 -u -50 -U -50 -m 100 -a AATGATACGGCGACCACCGAGATCTACACTCTTTCCCTACACGACGCTCTTCCGATCT -A GATCGGAAGAGCACACGTCTGAACTCCAGTCACNNNNNNATCTCGTATGCCGTCTTCTGC TTG’. For comparative analysis, 1,600,000 reads (ca 0.1× genome coverage of *N. furzeri*) were pseudorandomly subsampled from each biological replica of each species and resulting subsets were concatenated and analyzed together. The RepeatExplorer2 pipeline was run on the Galaxy server (The Galaxy Community 2022) with Metazoa version 3.0 protein domain database and automatic filtering of abundant repeats. In addition, the repeats were studied in each species independently, using a set of 7,125,000 reads (ca 0.5× coverage) and equivalent RepeatExplorer2 parameters. Calculation of G+C content and reciprocal BLAST were performed in GeneiousPrime (version 2020.1.2; https://www.geneious.com). To target potential centromeric repeats, the results of the single-species analysis were confined to high confidence satellites with estimated abundance in the genome at least 0.15% and monomer length <1kb only.

### 2.6 Identification of putative CENP-B box

SatDNA motifs with confirmed centromeric localization (see below) were manually inspected for the presence of 17-bp-long CENP-B box motif (Suntronpong et al. 2016) using Geneious Prime. Alignment of putative CENP-B sequences from human (*Homo sapiens*; Masumoto et al. 1989), threespine stickleback (*Gasterosteus aculeatus*; Cech and Peichel 2015), ninespine stickleback (*Pungitius pungitius;* Varadharajan et al. 2019), Asian swamp eel (*Monopterus albus;* Suntronpong et al. 2020) and turquoise killifish (*Nothobranchius fuzeri*; this work) were performed with a Geneious native algorithm with default settings for global alignment.

### 2.7 Fluorescence in situ hybridization (FISH)

#### 2.7.1. Preparation of FISH probes

We previously characterized Nfu-SatA and Nfu-SatB as the most abundant satellite DNA motifs in *N. furzeri* and *N. kadleci* (Štundlová et al. 2022). For FISH, the probe covering the whole monomer length 77 bp of Nfu-SatA was generated by 5’ labeling with Cy3 during synthesis (Generi Biotech, Hradec Králové, Czech Republic). In the same way, the probes for the Cl-36, Cl-127, Cl-260 and Cl-294, characterized for the first time in the present study (see below), have been prepared. In the case of Nfu-SatB (348-bp-long monomer), the clones with inserts containing three adjacent tandemly arrayed Nfu-SatB monomers which were prepared and verified in the previous study (Štundlová et al. 2022) were used for the FISH probe preparation. The entire plasmids were labeled by nick translation using a Cy3 NT Labeling Kit (Jena Bioscience, Jena, Germany). For the final probe mixture preparation, 250–500 ng of labeled plasmid and 12.5–25 μg of sonicated salmon sperm DNA (Sigma-Aldrich) were applied per slide. The final hybridization mixtures for each slide (15 μL) were prepared according to Sember et al. (2015).

#### 2.7.2. Standard FISH analysis

Single-color FISH experiments with Nfu-SatB probe were carried out following Sember et al. (2015) (slide pre-treatment, probe/chromosomes denaturation and hybridization conditions) and Yano et al. (2017) (post-hybridization washing), with modifications described in Štundlová et al. (2022). Briefly, following standard pre-treatment steps, chromosomes were denatured in 75% formamide in 2× SSC (pH 7.0) (Sigma-Aldrich) at 72 °C for 3 min. The hybridization mixture was denatured at 86 °C for 6 min. The hybridization took place overnight (17–24h) at 37 °C in a moist chamber. Subsequently, non-specific hybridization was removed twice in 1× SSC (pH 7.0) (65 °C, 5 min each) and once in 4× SSC in 0.01% Tween 20 (42 °C, 5 min), followed by washing in 1× PBS (1 min at room temperature; RT). Slides were dehydrated in an ethanol series (70%, 80%, and 96%, 2 min each) and then mounted in anti-fade containing 1.5 μg/mL DAPI (Cambio, Cambridge, UK).

#### 2.7.3. Non-denaturing FISH (ND-FISH)

Remaining five satDNA probes (Nfu-SatA, Cl-36, Cl-127, Cl-260 and Cl-294; more details provided in section 3.2) were mapped using ND-FISH according to Cuadrado and Jouve (2010) with some modifications. Briefly, a total of 30 µL of hybridization mixture containing 2 pmol/µL of oligonucleotides (labeled at 5’ end with Cy3) in 2× SSC were used per slide. Then the mixture was denatured at 80 °C for 5 min and immediately placed on ice. After that, the denatured hybridization mixture was transferred into the slides with neither pre-treatment steps nor chromosome denaturation. After two hours of hybridization at RT, the slides were washed with 4× SSC 0.2% Tween-20 at RT and shaking for 10 min, followed by 5 min washing in 4× SSC 0.1% Tween-20 also at RT and shaking. Chromosome preparations were then passed through ethanol series (70%, 80% and 96%, 3 min each) and then air dried. Chromosomes were counterstained with 20 µL of DABCO anti-fade (1,4-diazabicyclo(2.2.2)-octane containing 0.2 μg/mL DAPI (both Sigma-Aldrich) or in anti-fade containing 1.5 μg/mL DAPI Cambio, Cambridge, UK).

### 2.8. Microscopic analyses and image processing

Images from all cytogenetic methods were captured using a BX53 Olympus microscope equipped with an appropriate fluorescence filter set and coupled with a black and white CCD camera (DP30W Olympus). Images were acquired for each fluorescent dye separately using DP Manager imaging software (Olympus), which was further used also to superimpose the digital images with the pseudocolors (red for CMA_3_ and green for DAPI in case of fluorescence staining; blue for DAPI and red for Cy3 in case of FISH). Composite images were then optimized and arranged using Adobe Photoshop, version CS6.

At least 20 chromosome spreads per individual and method were analyzed. Chromosomes were classified according to Levan et al. (1964), but modified as m – metacentric, sm – submetacentric, st – subtelocentric, and a – acrocentric, where st and a chromosomes were scored together into st-a category.

## 3. Results

### 3.1 Cytogenetics

#### Basic karyotype characteristics

Individuals from all studied species displayed mostly the same 2n and highly similar proportion of chromosome categories as previously reported (Reichwald et al. 2009, 2015; Krysanov and Demidova 2018). The only exception was *N. ocellatus*, where we recorded 2n = 32 with the karyotype being composed exclusively of monoarmed (st-a) chromosomes, in contrast to previously reported 2n = 30 with one chromosome pair being large metacentric (Krysanov and Demidova 2018). The individuals studied by Krysanov and Demidova (2018) were later found to be members of a newly described closely related species *N. matanduensis* (Watters et al. 2020) (S. Simanovsky, pers. commun.). Finally, in line with the previous reports (Ewulonu et al. 1985; Krysanov and Demidova 2018), *Fundulosoma thierryi* and *N. guentheri* possessed male heterogametic X_1_X_1_X_2_X_2_/X_1_X_2_Y multiple sex chromosome system manifested by different chromosome counts between males and females (males had one chromosome less) and particularly in *N. guentheri* the male-limited neo-Y chromosome was discernible as the only large sm/st element in the complement.

## Distribution and composition of constitutive heterochromatin

Amount of constitutive heterochromatin varied among the studied *Nothobranchius* spp. (Supplementary Fig. 1). The outgroup taxon *F. thierryi* together with *Nothobranchius* species from the Southern clade possessed generally more heterochromatin segments than *N. ocellatus* and species from the Coastal clade where the narrow C-bands were confined mostly to the (peri)centromeric regions of overwhelming majority (*N. cardinalis, N. ocellatus*), two-thirds (*N. rubripinnis*), about half (*N. eggersi*, *N. guentheri*) or several chromosomes (*N. foerschi*) of the complement. Within the chromosome complements of *N. cardinalis*, *N. guentheri* and *N. rubripinnis*, the largest metacentric chromosome pair either lacked or had unremarkable/ notably smaller C-bands compared to the remainder of the chromosome set (Supplementary Fig. 1J–M). In *N. foerschi*, the largest metacentric chromosome pair possessed a distinct (peri)centromeric C-band, while the second largest metacentric pair displayed only tiny heterochromatin block (Supplementary Fig. 1I). By contrast, majority of large biarmed chromosomes in species of the Southern clade possessed large heterochromatin segments (see below). In addition to (peri)centromeric bands, heterochromatin accumulations were present on the short arms of several chromosomes in *N. eggersi*. In males of *N. guentheri*, neo-Y sex chromosome bore an apparent C-banded region on its long arms Supplementary Fig. 1J, arrowhead). The other species with known X_1_X_2_Y multiple sex chromosome system (*F. thierryi*) did not show any exceptional C-banding pattern on these sex chromosomes. Four st-a chromosomes in *F. thierryi* displayed remarkable heterochromatin blocks covering their short arms. In the Southern clade, *N. orthonotus* and *N. kuhntae* featured the highest amount and diversity of heterochromatin blocks which were distributed on multiple regions across the chromosome complement. This observation is consistent with large (peri)centromeric regions found previously in closely related *N. furzeri* and *N. kadleci* (Štundlová et al. 2022; see Supplementary Fig. 2A, B for comparison). On the other hand, chromosomes of *N. pienaari*, *N. krysanovi* and *N. rachovii* bore almost exclusively (peri)centromeric bands of variable lengths, some of them being remarkably large (Supplementary Fig. 1D–F). In the species with almost exclusively biarmed (metacentric or submetacentric) chromosomes and low 2n, namely *N. krysanovi* and *N. rachovii*, some (peri)centromeres were arranged as two large adjacent blocks. *N. krysanovi* also displayed additional interstitial heterochromatin blocks on several chromosomes. In *N. rachovii* only two large submetacentric chromosomes possessed very tiny interstitial bands in addition to (peri)centromeric ones.

Fluorescent staining revealed, besides few predominantly DAPI^+^ (AT-rich) bands (e.g., in *F. thierryi*, *N. orthonotus*), variable amount and distribution of CMA_3_ ^+^ (GC-rich) regions. Five species (*F. thierryi*, *N. pienaari*, *N. krysanovi* and *N. foerschi*) displayed just one pair of clear terminal or interstitial signals, highly likely overlapping with major ribosomal DNA (rDNA) cluster (cf. Sember et al. 2015 and references therein). Similar signals were revealed also on the neo-Y and at least one X chromosome of *N. guentheri* (Supplementary Fig. 3J). Several *N. guentheri* chromosomes also featured additional tiny centromeric signals on at least four chromosomes (Supplementary Fig. 3J, K). In *N. rachovii*, terminal CMA_3_ ^+^ signals were observed on the short arms of the smallest acrocentric chromosome pair, and at least four large metacentrics/submetacentrics had a tiny centromeric signal (Supplementary Fig. 3F). *N. ocellatus* and *N. eggersi* bore up to seven and up to four signals, respectively. *N. cardinalis* and *N. rubripinnis* shared the CMA_3_ pattern in the way that (peri)centromeres of all chromosomes were GC-rich except for the one pair of large metacentric chromosomes. Finally, almost all chromosome pairs in *N. orthonotus* and *N. kuhntae* had GC-rich (peri)centromeres, similarly to patterns found in *N. furzeri* and *N. kadleci* (Štundlová et al. 2022; Supplementary Fig. 2C, D).

### 3.2 Identification of candidate repeats

The comparative analysis of tandem repeats in representatives of the Southern (*N. furzeri*, *N. kadleci*, *N. orthonotus*, *N. rachovii*) and Coastal (*N. guentheri*, *N. rubripinnis*) clades revealed in total 21 high confidence satellites with various abundances (Table 2). The two most abundant tandem repeats, namely Cl-11 and Cl-26, were the previously studied putative centromeric repeats Nfu-SatB and Nfu-SatA, respectively. Besides *N. furzeri* and *N. kadleci*, these clusters were also enriched in *N. orthonotus*, however, limited or completely missing in *N. rachovii*, *N. guentherii* and *N. rubripinnis*, suggesting existence of different motifs in centromeres of these species. Interestingly, satellites Cl-36 and Cl-294 showed the opposite pattern, as they were present in *N. rubripinnis* and *N. guentherii* but missing in the rest of the surveyed taxa. Single-species analysis with more stringent criteria (estimated abundance in the genome at least 0.15%, monomer length <1kb) confirmed these results. Besides already identified markers, one more abundant satDNA (Cl-127) was identified to be shared by multiple species and was therefore included in further analysis, as well as Cl-260, which showed a high number of similarity hits with the above mentioned Nfu-SatA (Cl-26) (Table 3).

**Table 2.** High confidence satellites identified by comparative analysis with RepeatExplorer2

| Satellite | Monomer (bp) | GC (%) | Avg reads per replica |  |  |  |  |  | Notes |
| --- | --- | --- | --- | --- | --- | --- | --- | --- | --- |
|  |  |  | NFU | NKA | NOR | NRA | NGU | NRU |  |
| CI-11 | 348 | 41.1 | 2,933 | 1,508 | 643 | 50 | 23 | 9 | Previously identified Nfu-SatB* |
| CI-26 | 77 | 63.6 | 560 | 2,533 | 369 | 0 | 2 | 0 | Previously identified Nfu-SatA* |
| CI-28 | 169 | 36.1 | 466 | 164 | 4,700 | 34 | 18 | 50 |  |
| CI-36 | 48 | 45.8 | 0 | 0 | 0 | 0 | 195 | 2,107 | Specific for Coastal clade |
| CI-97 | 93 | 60.2 | 398 | 1,240 | 262 | 42 | 521 | 233 |  |
| CI-127 | 39 | 20.5 | 1,160 | 836 | 19 | 21 | 14 | 28 | High abundance in multiple species |
| CI-147 | 349 | 60.2 | 62 | 313 | 119 | 108 | 33 | 996 |  |
| CI-156 | 49 | 65.3 | 1,261 | 217 | 30 | 0 | 0 | 0 |  |
| CI-187 | 84 | 31 | 763 | 214 | 86 | 207 | 178 | 68 |  |
| CI-238 | 24 | 58.3 | 556 | 27 | 3 | 2 | 13 | 7 |  |
| CI-260 | 76 | 57.9 | 0 | 512 | 2 | 0 | 0 | 0 | Similarity hits with Nfu-SatA (CI-26) |
| CI-274 | 662 | 46.8 | 1 | 0 | 10 | 163 | 266 | 2 |  |
| CI-278 | 21 | 19 | 304 | 20 | 56 | 14 | 17 | 18 |  |
| CI-286 | 20 | 45 | 0 | 0 | 356 | 0 | 1 | 4 |  |
| **CI-294** | **63** | **31.7** | **0** | **0** | **0** | **0** | **193** | **137** | **Specific for Coastal clade** |
| CI-297 | 691 | 42.1 | 87 | 46 | 67 | 45 | 35 | 37 |  |
| CI-310 | 976 | 40.1 | 47 | 31 | 40 | 48 | 41 | 61 |  |
| CI-327 | 980 | 37.3 | 49 | 42 | 29 | 37 | 29 | 41 |  |
| CI-335 | 625 | 43.3 | 0 | 0 | 208 | 0 | 0 | 0 |  |
| CI-342 | 580 | 36.9 | 39 | 27 | 29 | 20 | 39 | 39 |  |
| CI-367 | 490 | 44.3 | 27 | 15 | 16 | 29 | 33 | 44 |  |

Markers selected for physical mapping are indicated in bold
\*Reichwald et al. 2009; Štundlová et al. 2022

**Table 3.**
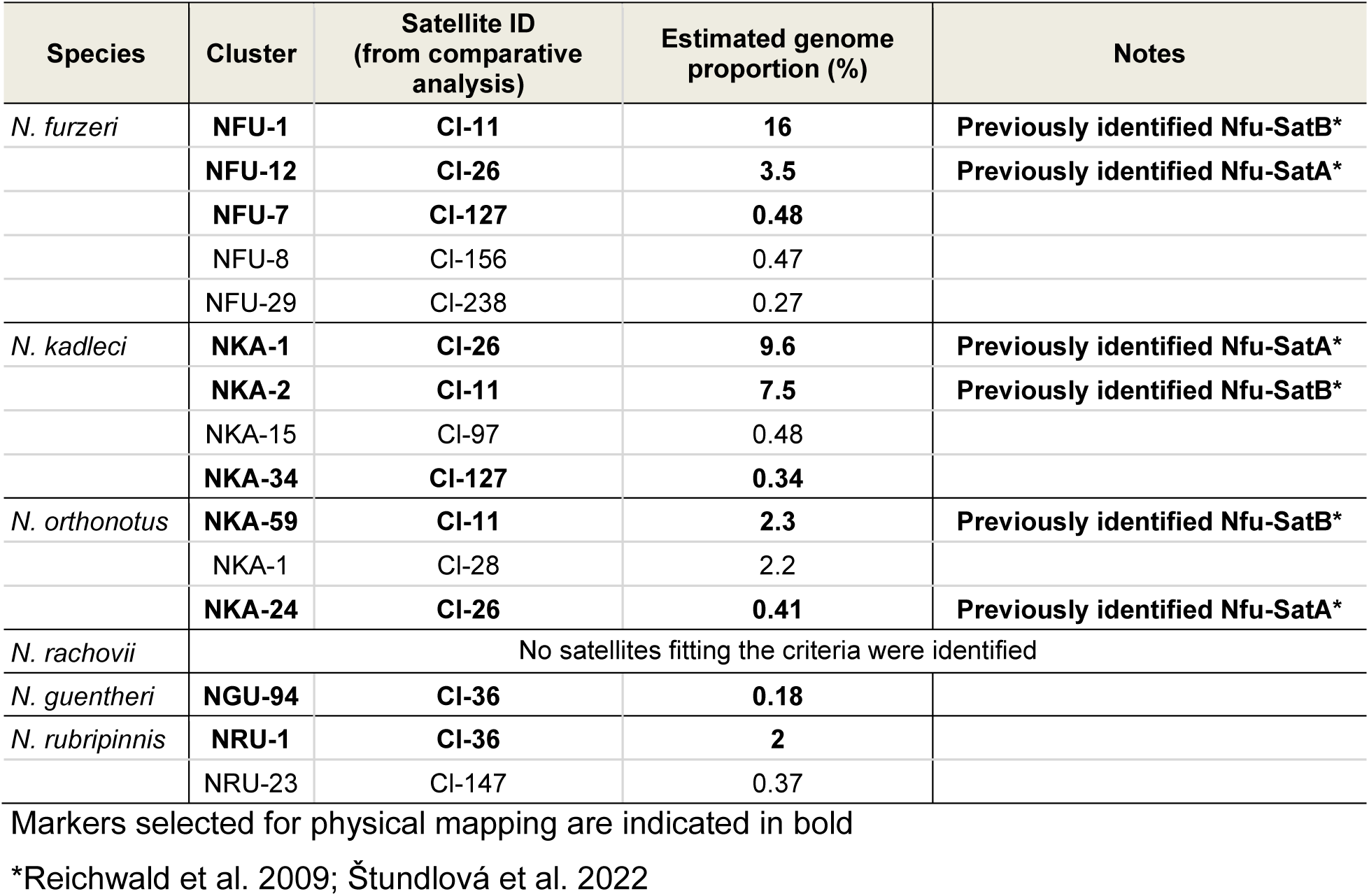
Species-specific analysis of the most abundant satellites (abundance >0.15% of the genome, monomer length < 1kb)

| Species | Cluster | Satellite ID<br>(from comparative<br>analysis) | Estimated genome<br>proportion (%) | Notes |
| --- | --- | --- | --- | --- |
| *N. furzeri* | **NFU-1** | **CI-11** | **16** | **Previously identified Nfu-SatB\*** |
|  | **NFU-12** | **CI-26** | **3.5** | **Previously identified Nfu-SatA\*** |
|  | **NFU-7** | **CI-127** | **0.48** |  |
|  | NFU-8 | CI-156 | 0.47 |  |
|  | NFU-29 | CI-238 | 0.27 |  |
| *N. kadleci* | **NKA-1** | **CI-26** | **9.6** | **Previously identified Nfu-SatA\*** |
|  | **NKA-2** | **CI-11** | **7.5** | **Previously identified Nfu-SatB\*** |
|  | NKA-15 | CI-97 | 0.48 |  |
|  | **NKA-34** | **CI-127** | **0.34** |  |
| *N. orthonotus* | **NKA-59** | **CI-11** | **2.3** | **Previously identified Nfu-SatB\*** |
|  | NKA-1 | CI-28 | 2.2 |  |
|  | **NKA-24** | **CI-26** | **0.41** | **Previously identified Nfu-SatA\*** |
| *N. rachovii* | No satellites fitting the criteria were identified |  |  |  |
| *N. guentheri* | **NGU-94** | **CI-36** | **0.18** |  |
| *N. rubripinnis* | **NRU-1** | **CI-36** | **2** |  |
|  | NRU-23 | CI-147 | 0.37 |  |

Markers selected for physical mapping are indicated in bold
\*Reichwald et al. 2009; Štundlová et al. 2022

Intriguingly, we found a putative CENP-B box sequence in Nfu-SatB repeat. Alignment to human CENP-B box sequence showed 0.47 identity, which is similar to other fish species (Supplementary Fig. 4). This finding along with the length of Nfu-SatB motif (348 bp; i.e. approx. twice the length of the nucleosome unit) may imply a possible role of Nfu-SatB in centromere function (Talbert and Henikoff 2020). However, no CENP-B box motif was recovered in other inspected repeats and particularly Cl-36 centromeric satellite of *N. rubripinis* is also too short (48 bp monomer) compared to other repeats with putative centromeric function.

### 3.3 Physical mapping of satDNA

FISH with Nfu-SatA (Cl-26) probe revealed detectable clusters only in *N. orthonotus* and *N. kuhntae* (Supplementary Fig. 5B–D). All signals were restricted to (peri)centromeric regions of almost all chromosomes, corroborating patterns found in *N. furzeri* and *N. kadleci* (Štundlová et al. 2022; Supplementary Fig. 2E, F). Whilst all *N. kuhntae* individuals shared the same pattern (i.e., all but one chromosome pair carrying the signal; Supplementary Fig. 5D), individuals of *N. orthonotus* displayed site-number variability, with the number of chromosomes lacking the signal being either four (1 male), five (1 male, 1 female), or six (two males, one female) chromosomes (Supplementary Fig. 5B–C).

Detectable clusters of Nfu-SatB (Cl-11) were found in (peri)centromeric regions of all chromosomes in *N. orthonotus*, *N. kuhntae* (i.e. the same pattern as in *N. furzeri* and *N. kadleci*; Štundlová et al. 2022 and Supplementary Fig. 2G, H), and in (peri)centromeric or terminal regions of about one-third of the chromosome complement in *N. pienaari* (Supplementary Fig. 6B–D). Besides these species of Southern clade, we also found clear hybridization patterns in three species of Coastal clade. Individuals of *N. eggersi* showed four signals placed terminally on short arms of st-a chromosomes. *N. foerschi* and *N. cardinalis* each carried one pair of st-a chromosomes with (peri)centromeric signals. The pair was small-sized in *N. cardinalis* and among the largest in *N. foerschi*. The Nfu-SatB loci in *N. foerschi* coincided with CMA_3_^+^ sites (compare Supplementary Figs 3I and 6I).

Satellite repeat Cl-36 was detected only in three species of Coastal clade: *N. rubripinnis* (from which it was isolated), *N. foerschi* and *N. guentheri* (Supplementary Fig. 7K, L, N). The repeat clusters were located exclusively in the (peri)centromeric regions but none of the mentioned species possessed them in all chromosomes. Studied *N. foerschi* and *N. guentheri* males displayed 12 and 16 signals, respectively (Supplementary Fig. 7K, L). In *N. rubripinnis*, 22 out of 36 chromosomes bore the signal (Supplementary Fig. 7N).

The second satellite limited to *N. rubripinnis* and *N. guentheri* (Cl-294) was hybridized in both these species, however, signals were detected only on the long arms of four chromosomes in *N. rubripinnis* (Supplementary Fig. 8A, B). The lack of signal in *N. guentheri* could be explained either by its abundance being below the FISH detection threshold, or by different organization of this repeat in the genome.

Cl-127, shared by *N. furzeri* and *N. kadleci,* was present in both sexes of these species, but no positive FISH signals were observed in *N. orthonotus* (Supplementary Fig. 8C–E). In both species, signals were localized in the long arm of two pairs of chromosomes in both males and females.

The last hybridized marker was Cl-260, bearing similarity hits with Nfu-SatA (Cl-26). Positive signals from this satDNA were observed in all centromeres in both sexes of *N. furzeri* and *N. kadleci*. The only difference in the signal pattern between these two species was related to additional prominent signals located terminally on the short arms of two (*N. furzeri*) and four (*N. kadleci*) chromosomes, respectively (Supplementary Fig. 8F, G).

**Fig. 1.**
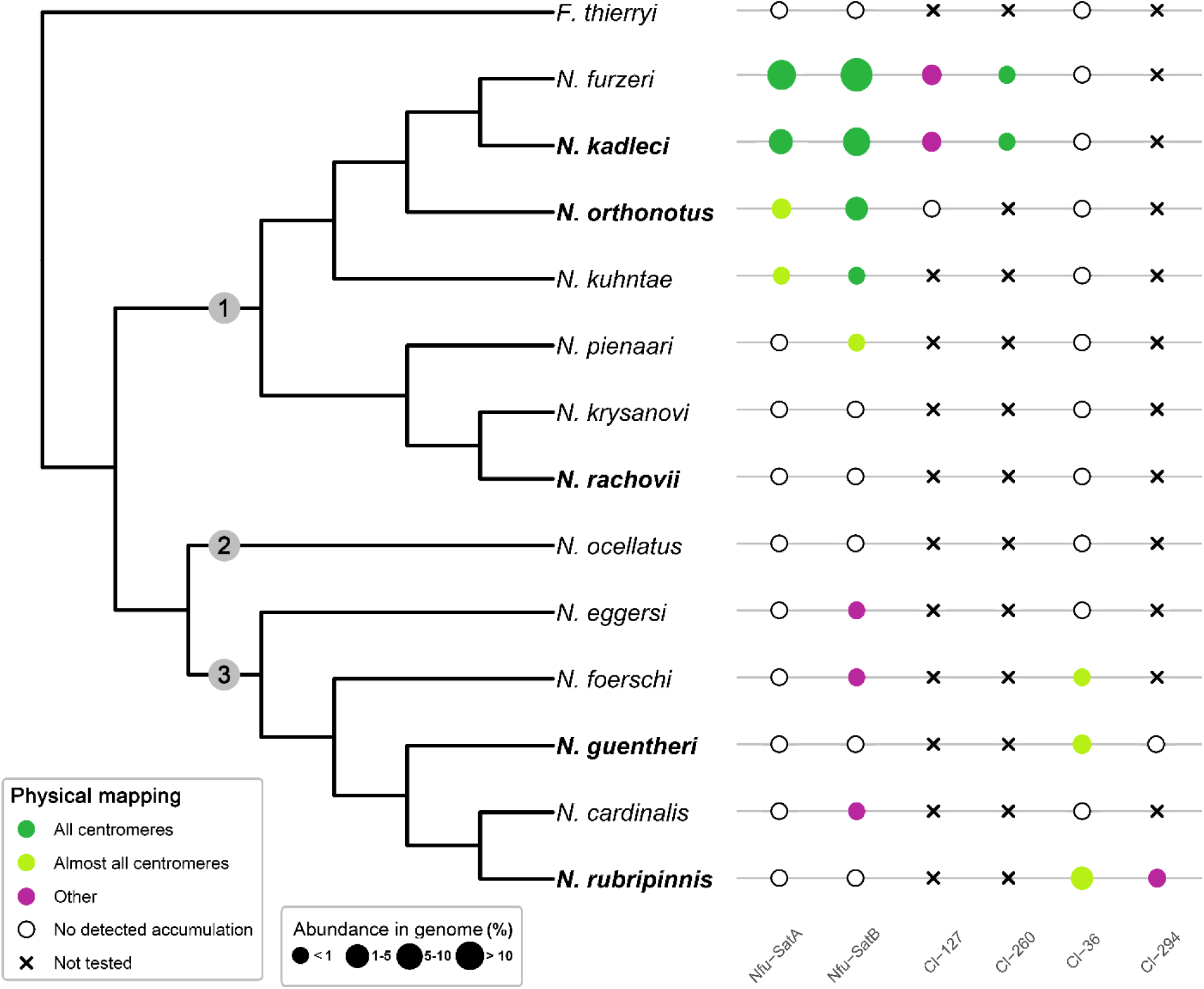
Phylogenetic relationships and patterns of selected repetitive DNA in inspected *Nothobranchius* species. Simplified phylogenetic tree is based on van der Merwe et al. (2021). Colored circles represent positive FISH signals in different chromosomal locations. The size of the circles reflects the abundance in the genome for respective satDNA. Abundance in the genome (%) is set as ranges. Lack of positive signals after FISH is demarcated by empty circles. Black crosses indicate that a given satDNA was not physically mapped in the particular species. Note that abundance in the genome might not perfectly correlate with chromosomal distribution revealed by physical mapping, because some portion of respective tandem repeats may be present in low-copy clusters undetectable by FISH. Species which were subject to RepeatExplorer2 analysis are shown in bold. Numbers in grey circles in the phylogenetic tree denote distinct *Nothobranchius* clades: 1) Southern Clade; 2) Ocellatus Clade; 3) Coastal Clade

## 4. Discussion

In the present study, we performed comparative cytogenetic and bioinformatic analyses of satellite DNA across the species of Southern and Coastal clade of the killifish genus *Nothobranchius* to reveal dynamics of repeats associated with centromeres and possible role of meiotic drive in their turnover.

Our results showed that the outgroup *F. thierryi* as well as *Nothobranchius* spp. of the Southern clade, namely *N. orthonotus*, *N. kuhntae*, *N. pienaari*, and *N. rachovii*, have in general more C-banded heterochromatin than representatives of the Coastal clade and their outgroup *N. ocellatus.* The presence of extended C-banded (peri)centromeric heterochromatin regions in large metacentric chromosomes of *N. pienaari*, *N. krysanovi* and *N. rachovii* (Supplementary Fig. 1D–F) is consistent with our previous findings in the remaining Southern-clade species, *N.* furzeri and *N. kadleci* (Štundlová et al. 2022 and Supplementary Fig. 2A, B) where we reported large amounts of (peri)centromeric heterochromatin in almost all chromosomes of the complement. By contrast, our present study shows only narrow C-bands in (peri)centromeric regions of majority of chromosomes in *N. cardinalis*, *N. ocellatus,* and *N. rubripinnis* (Supplementary Fig. 1G, L, M), and in about half-to-several chromosomes in *N. eggersi*, *N. foerschi* and *N. guentheri* (Supplementary Fig. 1H–J). Interestingly, with the sole exception of one chromosome pair in *N. foerschi*, large metacentric chromosomes originating from fusions either lacked or had unremarkable or notably smaller C-bands than other chromosomes in Coastal-clade species (*N. cardinalis*, *N. foerschi*, *N. guentheri* and *N. rubripinnis*). These findings indicate differences in mechanisms underpinning direction of karyotype change between Southern-clade and Coastal-clade killifishes.

To characterize the satDNA evolution, we sequenced and analyzed short reads of *N. guentheri*, *N. kadleci*, *N. orthonotus*, *N. rachovii* and *N. rubripinnis* together with data available for the model species *N. furzeri*. In total, the RepeatExplorer2 comparative analysis revealed 21 satellite sequences. We have analyzed the distribution of the most abundant satellites in all species (Nfu-SatA, Nfu-SatB, Cl-36 and Cl-127) and two additional markers (Cl-260 sharing similary with Nfu-SatA and Cl-294 specific for the Coastal clade) across both clades and the outgroups by FISH.

Štundlová et al. (2022) reported two satDNA motifs, Nfu-SatA and Nfu-SatB, previously identified in the *N. furzeri* strains (Reichwald et al. 2009, 2015) to be the most abundant repeat types in both the *N. furzeri* and *N. kadleci* genomes. Both Nfu-SatA and Nfu-SatB were mapped to (peri)centromeric constitutive heterochromatin blocks of varying sizes in these two sister species (see also Supplementary Fig. 2 C–F). Our results suggest that Nfu-SatA is restricted only to the *N. furzeri* lineage as it is present, although in lower abundance, also in (peri)centromeric regions of almost all chromosomes in *N. orthonotus* and *N. kuhntae* (Supplementary Fig. 5B–D). Lastly, the satellite Cl-260, was detected in the (peri)centromeric regions of all chromosomes in *N. furzeri* and *N. kadleci* only (Supplementary Fig. 8F, G). While Cl-260 is highly likely a new sequence variant of Nfu-SatA, specific for *N. kadleci* genome only (Table 2), its high sequence similarity with Nfu-SatA was apparently responsible for observing a positive hybridization also in (peri)centromeres of *N. furzeri*.

Nfu-SatB was also detected in (peri)centromeric regions of all chromosomes in *N. orthonotus* and *N. kuhntae* (Supplementary Fig. 6B, C). However, it was further present in detectable amounts also in *N. pienaari* as well as *N. eggersi*, *N. foerschi*, and *N. cardinalis* of the Coastal clade (Supplementary Fig. 6D, H, I, K). The Nfu-SatB signals were located terminally on the short arms of two chromosome pairs in *N. eggersi*, while they resided in (peri)centromeric regions of about one-third of the complement in *N. pienaari* and one chromosome pair of each *N. foerschi* and *N. cardinalis*. Observed pattern is consistent with the “library” hypothesis (Fry and Salser 1977; Ruiz-Ruano et al. 2016) as Nfu-SatB seems to be shared across *Nothobranchius* spp. but it got amplified and associated with centromeres in the *N. furzeri* lineage of the Southern clade.

While Nfu-SatA and Nfu-SatB were found restricted to Southern clade species, satellites Cl-36 and Cl-294 mirrored this pattern as they were detected in the Coastal clade only. The Cl-294 was localized on the long arms of only four chromosomes in *N. rubripinnis* and could not have been detected by FISH on chromosomes of *N. guentheri* (Supplementary Fig. 8A, B). However, Cl-36 was successfully mapped in three representatives of the Coastal clade (Supplementary Fig. 7K, L, N). The hybridization signals were detected exclusively in the (peri)centromeric regions of majority but not all chromosomes in *N. rubripinnis, N. foerschi*, and *N. guentheri.* Corroborating the C- and fluorescent-banding patterns, Cl-36 clusters were absent in (peri)centromeres of some large metacentric chromosomes (Supplementary Fig. 7K, N). This observation is analogous to previously reported satellite-free centromeres, which emerged upon Robertsonian fusions in zebras (Cappelletti et al. 2022).

Interestingly, none of the above tested markers was detected in centromeres of *N. rachovii* (Supplementary Figs. 3F, 5F, 6F, 7H) and the RepeatExplorer2 analysis failed to identify any potentially centromeric satellites in this species (Tab. 2, Tab. 3). Since blocks of (peri)centromeric heterochromatin (Supplementary Fig. 1F), visible on most *N. rachovii* chromosomes, suggest the presence of tandem arrays rather than satellite-free centromeres, a possible explanation might be that microsatellites are the involved sequences in this case. Centromeric localization of short repeat motifs has been described before in various organisms (e.g. Kim et al. 2002; Chang et al. 2008) and their presence could escape RepeatExplorer2 analysis as this tool is known to omit low complexity sequences (Novák et al. 2020).

Furthermore, a putative CENP-B box motif was identified in Nfu-SatB centromeric satellite. As in other fish species, its sequence showed overall 0.47 identity to the human CENP-B box (Supplementary Fig. 4). However, different nucleotide positions are conserved in various fishes (Cech and Peichel 2015; Varadharajan et al. 2019; Suntronpong et al. 2020). In addition, the CENP-B motif was not recovered in any other killifish (peri)centromeric satellites, including Cl-36. Thus, further research is needed to assess the contribution of Nfu-SatB-linked CENP-B box-like motif to centromere function. Differences between X- and Y-linked (peri)centromeric heterochromatin comprising Nfu-SatA and Nfu-SatB were reported in *N. furzeri* and *N. kadleci*, with the Y-linked heterochromatin being considerably reduced (Štundlová et al. 2022). It was hypothesized that this is due to absence of centromere drive on the Y chromosome as it is never transmitted via female meiosis (cf. Yoshida and Kitano 2012; Pokorná et al. 2014). Identification of putative centromeric repeat in the Coastal clade potentially presents an opportunity to test this hypothesis as *N. guentheri* has a multiple sex chromosome system of the X_1_X_2_Y type, in which neo-Y and one of the X chromosomes can be identified by CMA_3_ staining (Supplementary Fig. 3J). However, FISH with Cl-36 failed to detect any satDNA clusters on both the neo-Y and the CMA_3_-positive X chromosome.

It was hypothesized that karyotype evolution is driven by meiotic drive in many animal lineages (Pardo-Manuel de Villena and Sapienza 2001; Blackmon et al. 2019), including fishes (Yoshida and Kitano 2012; Molina et al. 2014), particularly by a nonrandom segregation of rearranged chromosome in female meiosis of heterokaryotypes, due to inherent asymmetry of female meiosis and polarity of a meiotic spindle. Stronger spindles should bind bigger centromeres (Chmátal et al. 2014; Akera et al. 2019; Kursel and Malik 2018; Kumon and Lampson 2022). Yet the direction of the nonrandom segregation is not set in stone. Reversals of spindle polarity supposedly occurred in many phylogenetic groups, which could explain differences in trends in karyotype evolution between related taxa (Pardo-Manuel de Villena and Sapienza 2001; Yoshida and Kitano 2012; Blackmon et al. 2019). It is tempting to speculate that in *N. furzeri* and *N. kadleci*, the egg has a stronger spindle pole than in the other species under study, as they have considerably larger (peri)centromeric heterochromatin blocks comprising Nfu-SatA and Nfu-SatB in all chromosomes but the Y chromosome. Interestingly, in both *N. furzeri* and *N. kadleci* the large blocks of (peri)centric heterochromatin coincide with higher numbers of chromosome arms, but not with different number of chromosomes than expected when compared to karyotypes of other *Nothobranchius* spp. (Krysanov and Demidova 2018). It suggests that evolution of satellite DNA in *Nothobranchius* species is associated either with intrachromosomal rearrangements or centromere repositioning, i.e. inactivation of an existing centromere and de novo formation of a new one elsewhere on the chromosome (cf. Amor et al. 2004; Cappelletti et al. 2022).

To conclude, our cytogenetic and bioinformatic data suggests that centromere drive operates in *Nothobranchius* killifishes and shape their karyotypes. Reversal of spindle polarity probably occurred in the Southern clade and changed direction of the drive. Further research is needed to parse causes and consequences of karyotype evolution in the killifishes.

## Supporting information

Supporting material for Volenikova et al.

## Acknowledgements

We would like to thank B. Nagy, A. Nikiforov and H. Hengstler for providing part of the study material and A. Nikiforov for his help in breeding and keeping fishes. We are also grateful to P. Šejnohová for the laboratory assistance.

## Authors’ contributions

Conceptualization: AS, PN; Data curation: PN, AV, PM, MA; Formal analysis: PN, AV; Funding acquisition: AS; Investigation: AV, KL, PM, TP, MA, JŠ, ŠP, SAS, MJ, PN, AS; Methodology: AS, PN, AV, PM; Project administration: AS, PN; Resources: AS, PN, MR; Supervision: AS, PN, MR; Validation: AV, PM, AS, PN, MA; Writing original draft: AV, AS, PN, PM; Writing—review & editing: PN, AS, AV, PM, MA, JŠ, MR, SAS.

## Availability of data and material

The following nucleotide sequences were deposited in Sequence Read Archive (SRA): XXX. All other relevant data are within the paper and its Supporting Information file.

## Declarations

### Ethics approval

To prevent fish suffering, all handling of fish individuals followed European standards in agreement with §17 of the Act No. 246/1992 Coll. The procedures involving fishes were supervised by the Institutional Animal Care and Use Committee of the Institute of Animal Physiology and Genetics CAS, v.v.i., and the supervisor’s permit number CZ 02361 was certified and issued by the Ministry of Agriculture of the Czech Republic. The experiments with *N. foerschi* and *N. cardinalis* were approved by the Ethics Committee of Severtsov Institute of Ecology and Evolution (Order No. 27 of November 9, 2018). For direct preparations of chromosomes from the kidney, fishes were euthanized using 2-phenoxyethanol (Sigma-Aldrich) before organ sampling. Fin samples (a narrow strip of the caudal fin) were taken from live individuals after fishes were anesthetized using MS-222 (Merck KGaA, Darmstadt, Germany).

## Abbreviations

2n: diploid chromosome number
a: acrocentric chromosome
CMA_3_: Chromomycin A_3_
DABCO: 1,4-diazabicyclo(2.2.2)-octane
DAPI: 4’,6-diamidino-2-phenylindole
FISH: fluorescence *in situ* hybridization
gDNA: genomic DNA
m: metacentric chromosome
ND-FISH: non-denaturing FISH
p-arm: short chromosome arm
PBS: phosphate-buffered saline
q-arm: long chromosome arm
rDNA: ribosomal DNA
RT: room temperature
satDNA: satellite DNA
SRA: Sequence Read Archive
SSC: saline-sodium citrate
st: subtelocentric chromosome

