## Supporting material for Volenikova et al. for "Fast centromeric repeat turnover provides a glimpse into satellite DNA evolution in *Nothobranchius* annual killifishes"

**The PDF file includes:**

Supplementary Fig. 1
Supplementary Fig. 2
Supplementary Fig. 3
Supplementary Fig. 4
Supplementary Fig. 5
Supplementary Fig. 6
Supplementary Fig. 7
Supplementary Fig. 8
Supplementary Table. 1

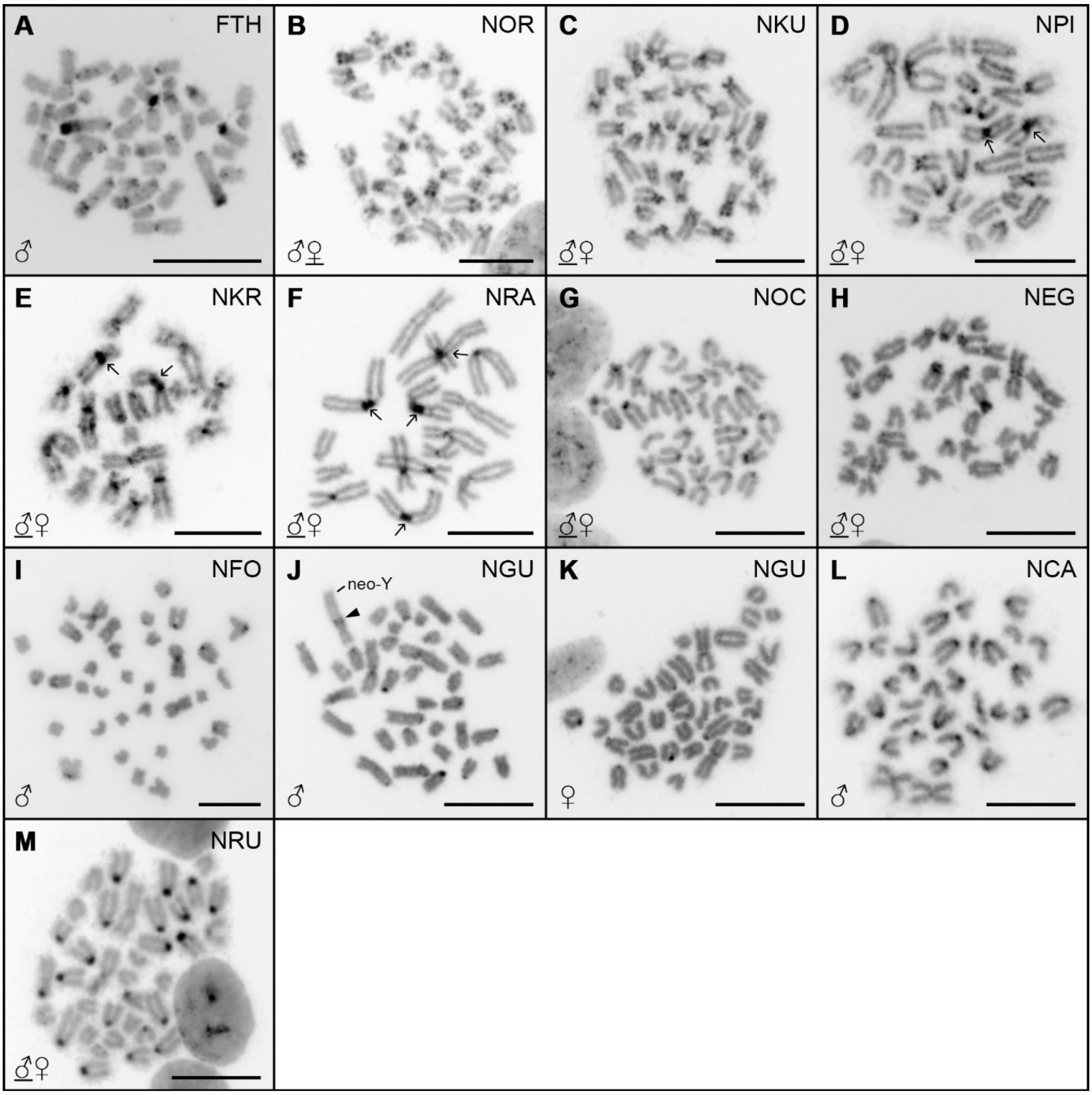


**Supplementary Fig. 1** Mitotic metaphases of *F. thierryi* and *Nothobranchius* spp. after C-banding. Sex of the studied individuals is indicated and eventually underlined where both sexes (if studied) presented the same distribution pattern (i.e. except for *N. guentheri*; J, K). Arrows indicate examples of huge (peri)centromeric heterochromatin blocks in expected fusion sites on large metacentric chromosomes in the Southern-clade species (D–F). Neo-Y chromosome in *N. guentheri* male (J) is identified based on its distinctive morphology; full arrowhead points to the heterochromatin block representing an assumed fusion site. Chromosomes stained with DAPI (inverted colors). Scale bar = 10 μm

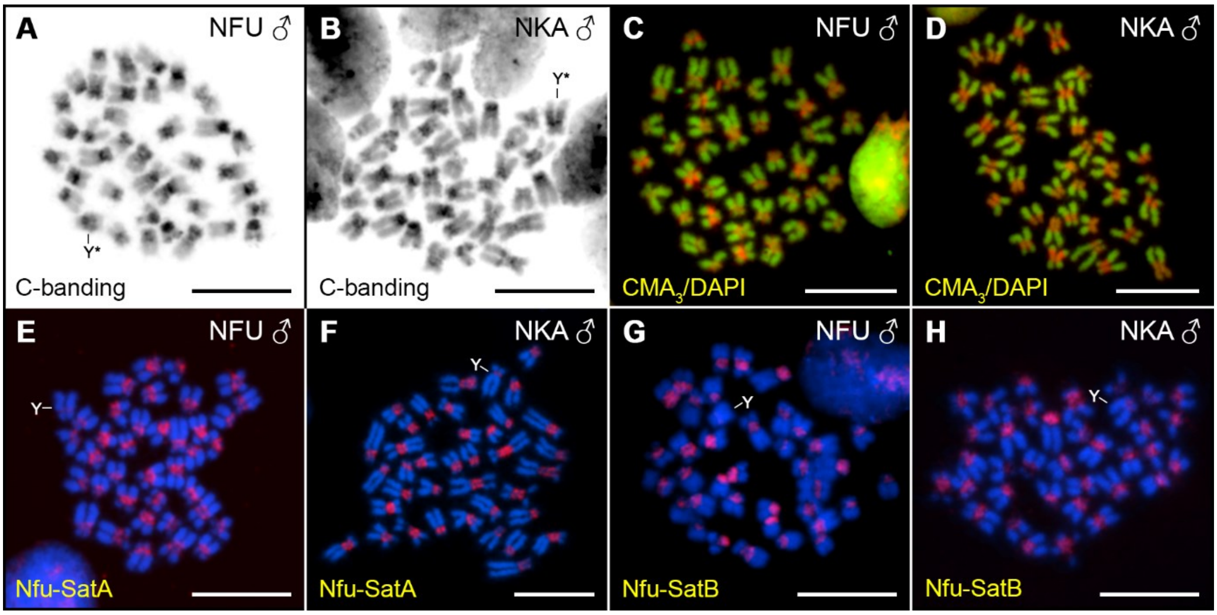


**Supplementary Fig. 2** Mitotic metaphases of *N. furzeri* and *N. kadleci* after various cytogenetic treatments. (A, B) C-banding, (C, D), $CMA_3$/DAPI staining, (E, F) FISH with Nfu-SatA probe, (G, H), FISH with Nfu-SatB probe. All results originate from the experiments published in our previous work (Štundlová et al. 2022) and serve here as a direct comparison of patterns revealed in the present study. The presented metaphases belong to male representatives of the same populations as used in the present study: *N. furzeri* MZCS-222 and *N. kadleci* MZCS-91. Assumed Y sex chromosomes are marked if detectable. Chromosomes were counterstained with DAPI (blue). Images follow the same color coding as the remaining supplementary figs in the present study (depending on the method used). Scale bar = 10 μm

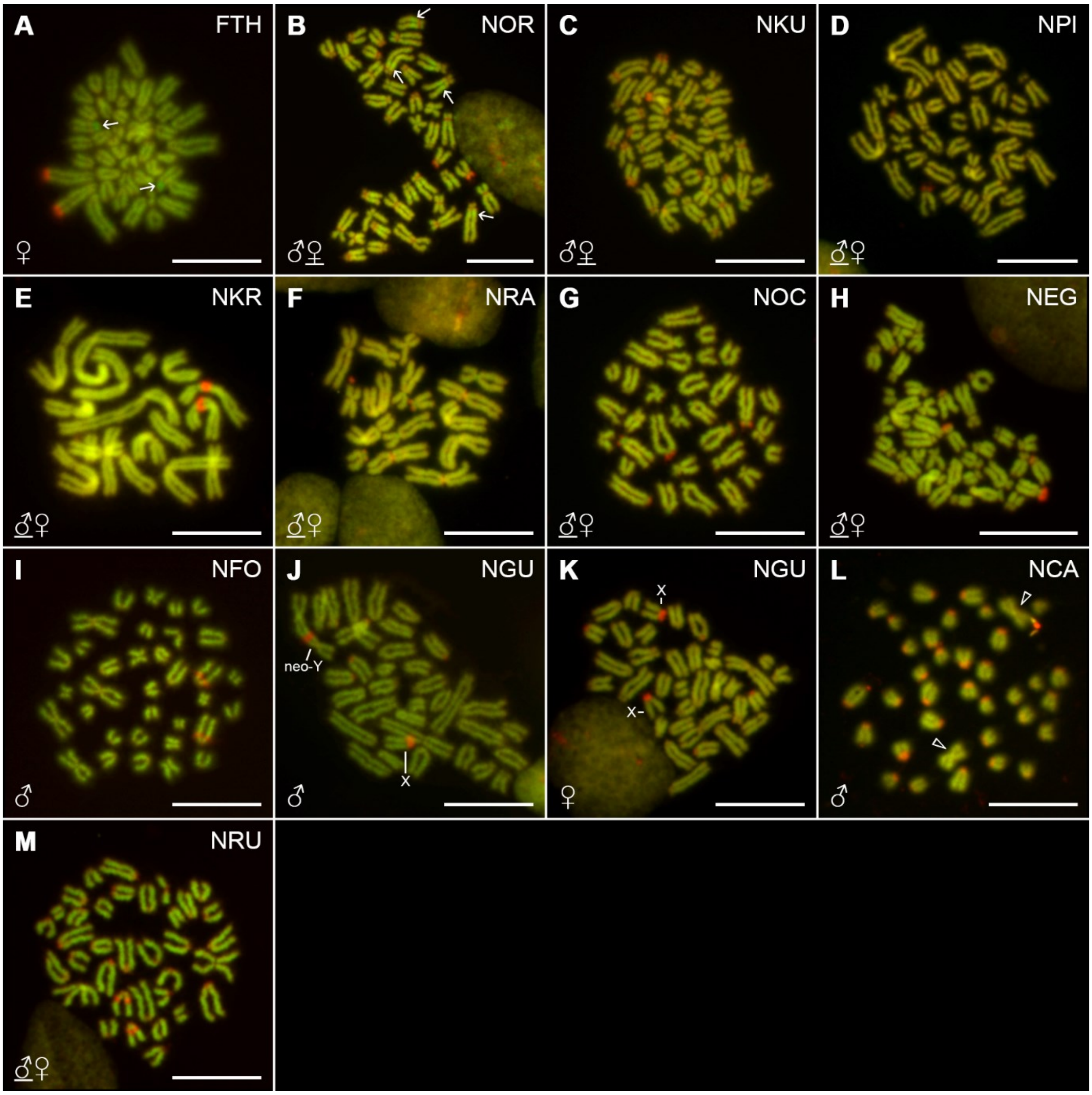


**Supplementary Fig. 3** Mitotic metaphases of *F. thierryi* and *Nothobranchius* spp. after $CMA_3$/DAPI staining. Sex of the studied individuals is indicated and eventually underlined where both sexes (if studied) presented the same distribution pattern (i.e. except for *N. guentheri*; J, K). For better contrast, images were pseudocolored in red (for $CMA_3$) and green (for DAPI). Neo-Y chromosome and one of the X chromosomes in *N. guentheri* male (J) are identified based on distinctive morphology and shared strong $CMA_3^+$ signals, respectively. Both X homologs are marked in *N. guentheri* female (K). In *N. cardinalis* (L) empty arrowheads point to the only metacentric chromosomes in the complement which are also the only elements lacking the (peri)centromeric $CMA_3$ signals. Arrows point to examples of rarely observed pronounced AT-rich regions (A, B). Scale bar = 10 μm

```
                  *  *  *  *              *        *  *  *  *     identity
H. sapiens     C  T  T  C  G  T  T  G  G  A  A  A  C  G  G  G  A
G. aculeatus   A  A  A  G  G  T  T  G  G  A  A  A  A  C  T  A  T     0.47
P. pungitus    T  A  G  T  T  T  T  A  G  A  A  C  C  T  G  A  N     0.44
M. albus       A  T  A  A  C  A  C  T  G  A  A  A  T  G  G  T  T     0.41
N. furzeri     C  T  T  C  G  T  G  C  A  A  C  A  T  A  A  T  T     0.47
```

**Supplementary Fig. 4** A putative CENP-B box motif in Nfu-SatB satellite in the turquoise killifish, *Nothobranchius furzeri*. Alignment shows 0.47 identity to the human CENP-B box sequence, which is comparable to those characterized in threespine stickleback (*Gasterosteus aculeatus*; 0.47 identity), ninespine stickleback (*Pungitus pungitus;* 0.44 identity) and Asian swamp eel (*Monopterus albus;* 0.41 identity). Conserved nucleotides of the human CENP-B box motif are indicated in red and functionally important domains for primates (Suntronpong et al. 2016; see References in the main MS body) are marked by asterisks

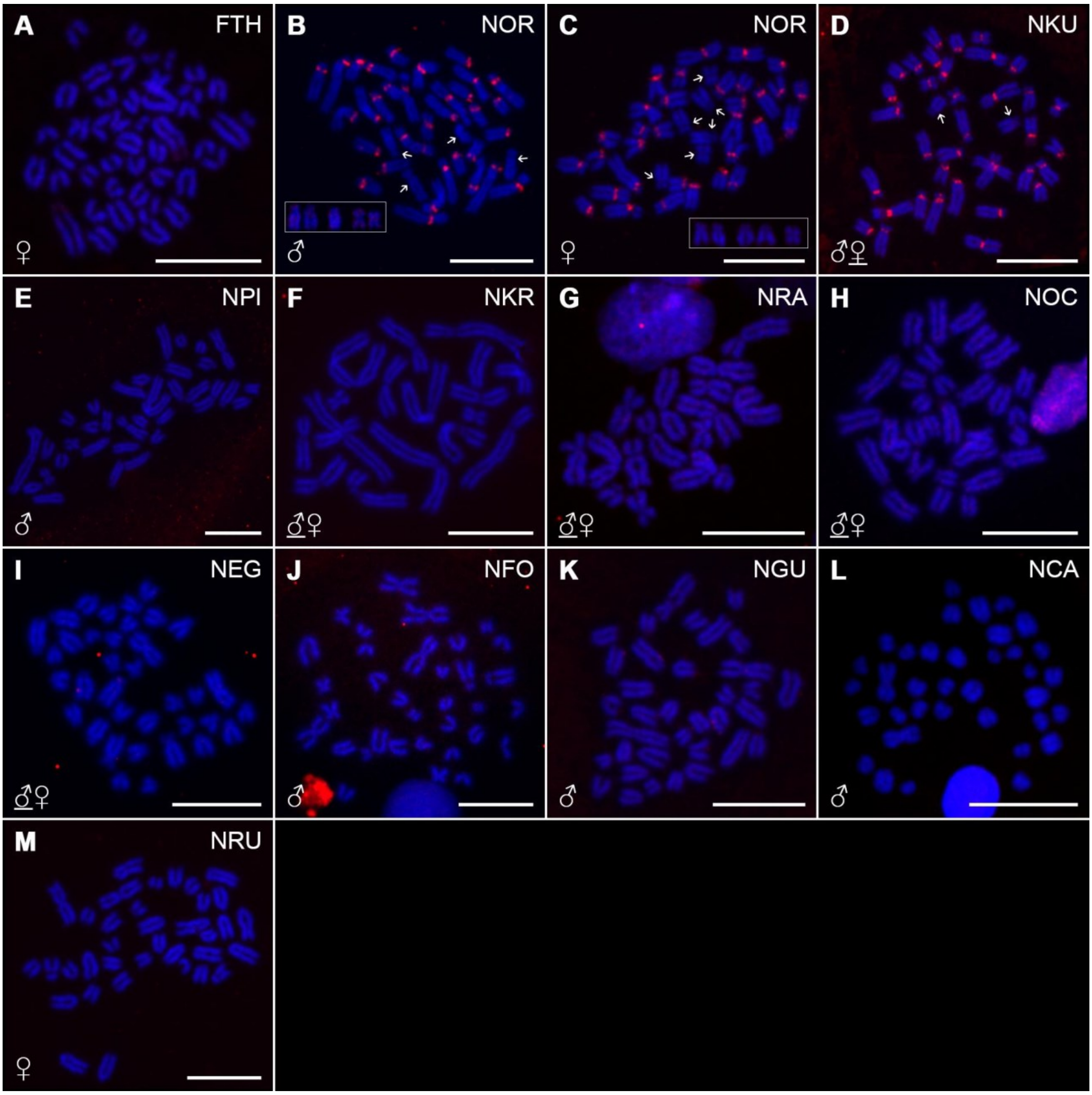


**Supplementary Fig. 5** Mitotic metaphases of *F. thierryi* and *Nothobranchius* spp. after FISH with Nfu-SatA repeat (red signals). Sex of the studied individuals is indicated and eventually underlined where both sexes (if studied) presented the same distribution pattern (i.e. except for *N. orthonotus*; B, C). In *N. orthonotus* (B, C), arrows point to chromosomes lacking the (peri)centromeric signals. Polymorphic patterns regarding this feature are framed. Chromosomes were counterstained with DAPI (blue). Scale bar = 10 μm

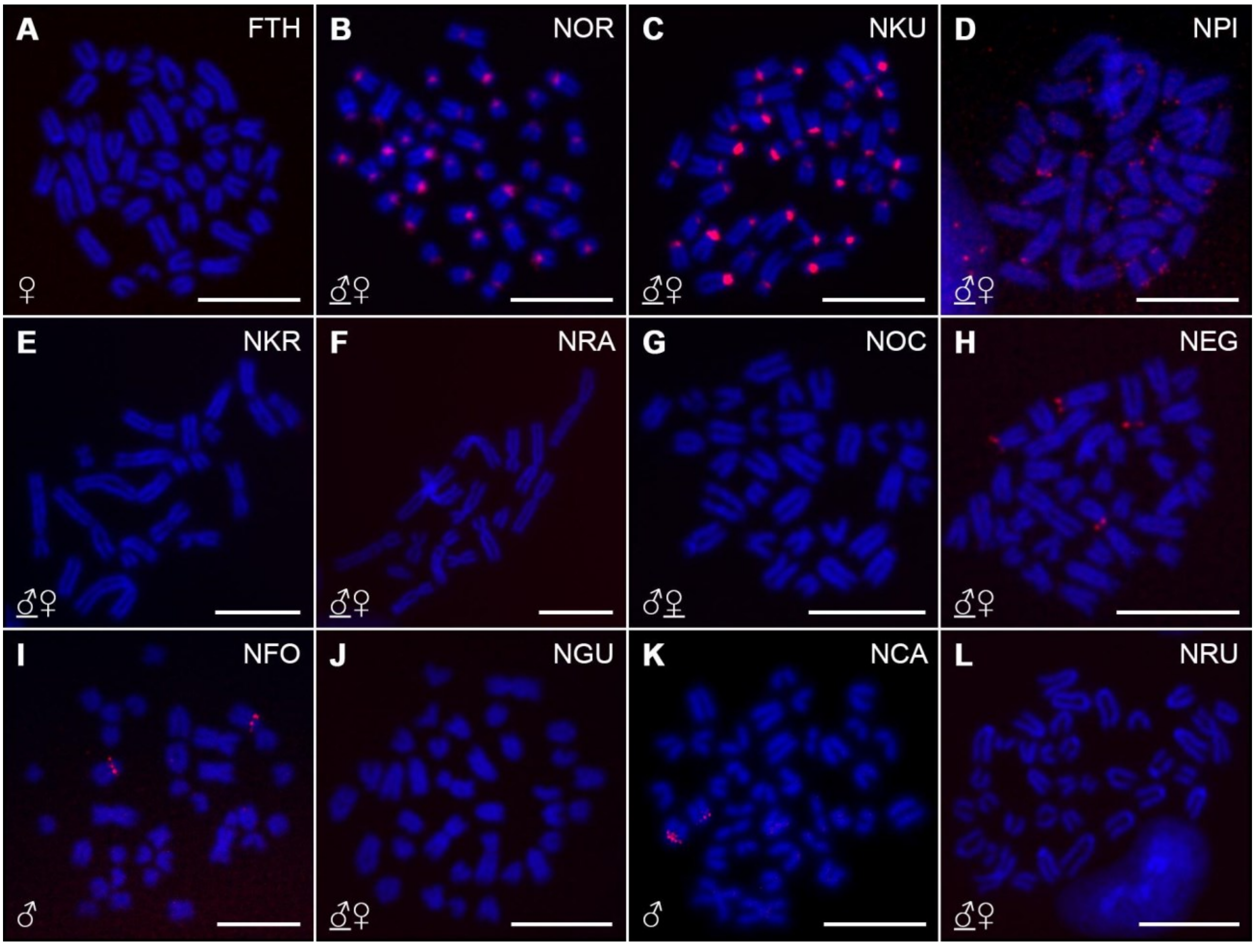


**Supplementary Fig. 6** Mitotic metaphases of *F. thierryi* and *Nothobranchius* spp. after FISH with Nfu-SatB repeat (red signals). Sex of the studied individuals is indicated and eventually underlined where both sexes (if studied) presented the same distribution pattern. Chromosomes were counterstained with DAPI (blue). Scale bar = 10 μm

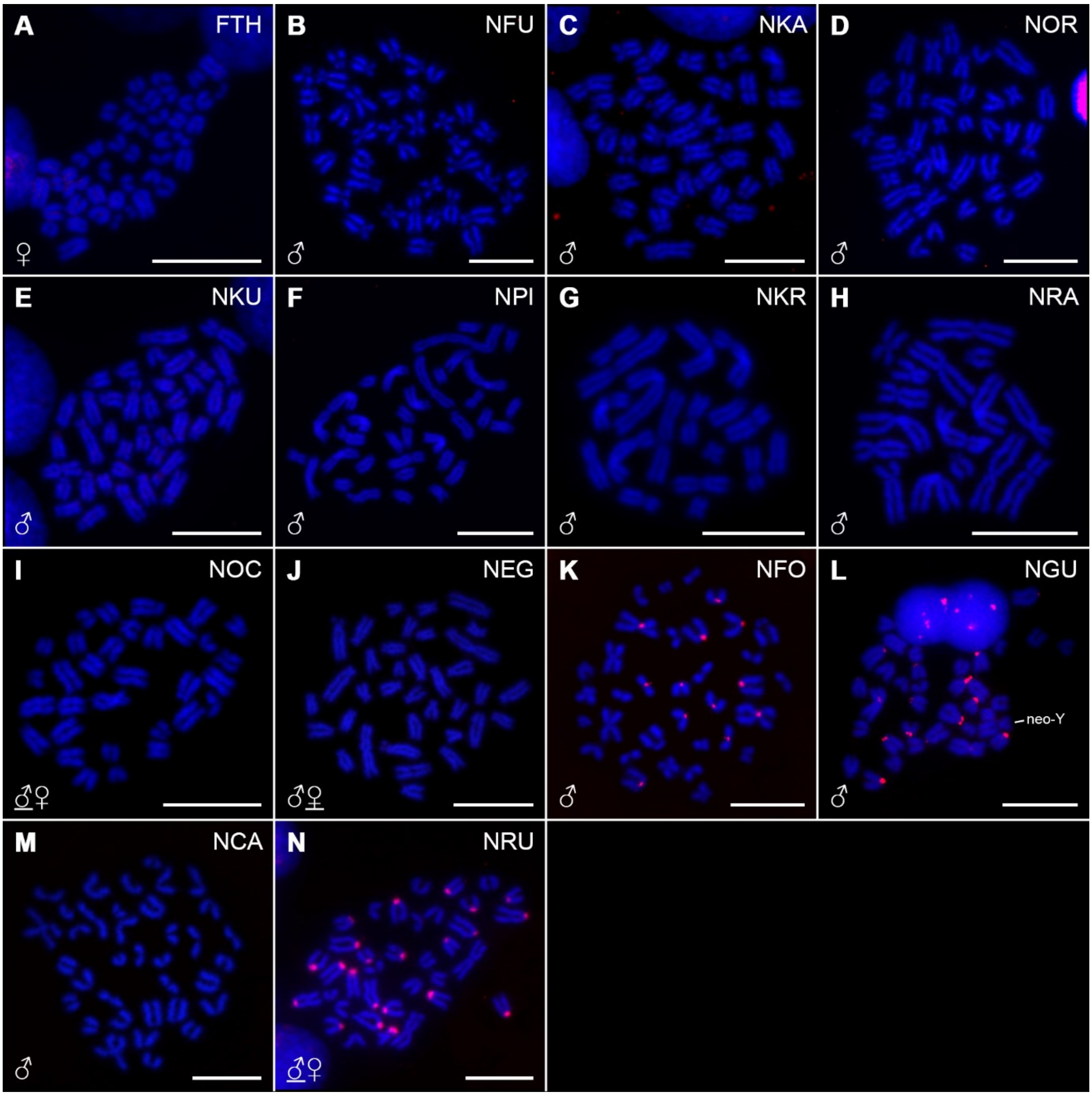


**Supplementary Fig. 7** Mitotic metaphases of *F. thierryi* and *Nothobranchius* spp. after FISH with Cl-36 repeat (red signals). Sex of the studied individuals is indicated and eventually underlined where both sexes (if studied) presented the same distribution pattern. Neo-Y chromosome in *N. guentheri* male (L) is identified based on distinctive morphology. Chromosomes were counterstained with DAPI (blue). Scale bar = 10 μm

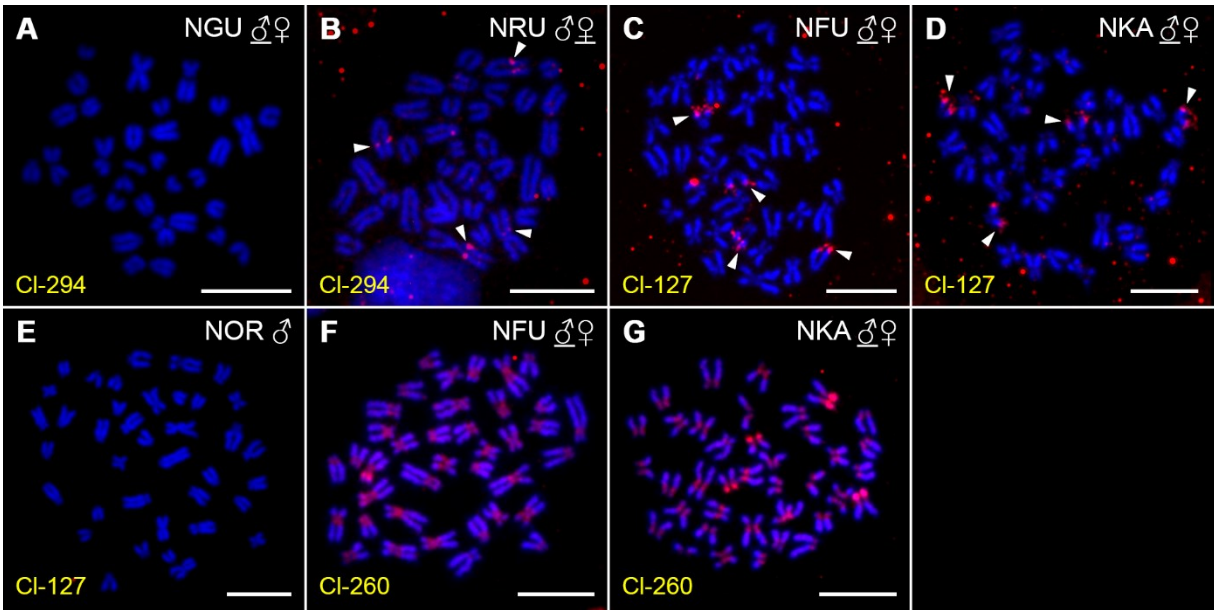


**Supplementary Fig. 8** Mitotic metaphases of selected *Nothobranchius* spp. after FISH with three different satDNA probes (red signals and arrowheads in B–D). Sex of the studied individuals is indicated and eventually underlined where both sexes (if studied) presented the same distribution pattern. Chromosomes were counterstained with DAPI (blue). Scale bar = 10 µm

**Supplementary Table 1** A detailed list of studied *Nothobranchius* killifish species with their sample sizes (N) used for each method, population/collection codes, source/geographic origin and GPS coordinates of sampling localities

| Clade | Species | Code | *N* | | | | | | | | | Population (collection code) | Source / locality | GPS coordinates |
|---|---|---|---|---|---|---|---|---|---|---|---|---|---|---|
| | | | C-banding | $CMA_3$ | Nfu-SatA | Nfu-SatB | Cl-127 | Cl-260 | Cl-36 | Cl-294 | Summary | | | |
| outgroup | *Fundulosoma thierryi* Ahl, 1924 | FTH | 1♂, 2♀ | 2♀ | 2♀ | 2♀ | – | – | 3♀ | – | 1♂, 3♀ | aquarium strain | – | – |
| Southern clade | *Nothobranchius furzeri* Jubb, 1971 | NFU | * | * | * | * | 1♂, 1♀ | 1♂, 1♀ | 2♂ | – | 1♂, 1♀ | MZCS-222 | Chefu, Mozambique | 21°52'24.8"S 32°48'2.3"E |
| | *N. kadleci* Reichard, 2010 | NKA | * | * | * | * | 1♂, 1♀ | 1♂, 1♀ | 1♂ | – | 1♂, 1♀ | MZCS-91 | Gorongosa, Mozambique | 20°41'16.6"S 34°6'21.9"E |
| | *N. orthonotus* (Peters, 1844) | NOR | 2♂, 3♀ | 1♂, 1♀ | 3♂, 3♀ | 1♂, 1♀ | 1♂, 2♀ | – | 1♂ | – | 3♂, 3♀ | MZCS-02 | Limpopo, Mozambique | 24°03'48.5"S 32°43'55.9"E |
| | *N. kuhntae* (Ahl, 1926) | NKU | 4♂, 1♀ | 1♂, 2♀ | 3♂, 3♀ | 1♂, 1♀ | – | – | 2♂ | – | 4♂, 3♀ | MZCS-528 | Pungwe, Mozambique | 19°41'50.5"S 34°46'58.6"E |
| | *N. pienaari* Shidlovskyi, Watters & Wildekamp, 2010 | NPI | 2♂, 2♀ | 2♂, 3♀ | 2♂ | 2♂, 2♀ | – | – | 1♂ | – | 2♂, 3♀ | MZCS-505 | Limpopo, Mozambique | 23°31'47.2"S 32°34'40.6"E |
| | *N. krysanovi* Shidlovskyi, Watters & Wildekamp, 2010 | NKR | 2♂, 2♀ | 1♂, 2♀ | 1♂, 2♀ | 1♂, 2♀ | – | – | 1♂ | – | 2♂, 2♀ | aquarium strain; MZCS-249 | Quelimane, Mozambique | 17°48'52.2"S 36°54'49.4"E |
| | *N. rachovii* Ahl, 1926 | NRA | 2♂, 2♀ | 1♂, 1♀ | 1♂, 1♀ | 1♂, 1♀ | – | – | 1♂ | – | 2♂, 2♀ | MZCS-096 | Beira Airport, Mozambique | 19°48'48.8"S 34°54'17.6"E |
| Ocellatus clade | *N. ocellatus* (Seegers, 1985) | NOC | 1♂, 1♀ | 1♂, 1♀ | 1♂, 1♀ | 1♂, 1♀ | – | – | 1♂, 1♀ | – | 1♂, 1♀ | – | Nyamwage, Tanzania | – |
| Coastal clade | *N. eggersi* Seegers, 1982 | NEG | 2♂, 1♀ | 1♂, 1♀ | 2♂, 1♀ | 1♂, 1♀ | – | – | 2♂, 1♀ | – | 2♂, 1♀ | T52 | Bagamoyo, Tanzania | 6°28'55.9"S 38°54'51.5"E |
| | *N. foerschi* Wildekamp & Berkenkamp, 1979 | NFO | 2♂ | 2♂ | 2♂ | 2♂ | – | – | 2♂ | – | 2♂ | CI 57 | Soga, Tanzania | 6°50'13.2"S 38°50'45.6"E |
| | *N. guentheri* (Pfeffer, 1983) | NGU | 3♂, 3♀ | 4♂, 2♀ | 1♂, 2♀ | 1♂, 2♀ | – | – | 2♂ | 2♂, 1♀ | 4♂, 3♀ | aquarium strain | Zanzibar, Tanzania | – |
| | *N. cardinalis* Watters, Cooper & Wildekamp, 2008 | NCA | 1♂ | 1♂ | 1♂ | 1♂ | – | – | 1♂ | – | 1♂ | TTKSN 17-12 | Matandu, Tanzania | 9°30'04.0"S 38°13'49.0"E |
| | *N. rubripinnis* Seegers, 1986 | NRU | 1♂, 1♀ | 1♂, 2♀ | 2♀ | 2♂ | – | – | 2♂, 2♀ | 1♂, 2♀ | 2♂, 2♀ | T33 | Kitonga, Tanzania | 7°12'40.5"S 39°10'30.9"E |

* reported in Štundlová et al. 2022 (see References in the main MS body)
